# A conserved matrix-centred core links fibrosis across human organs and mouse models

**DOI:** 10.64898/2026.11.13.766356

**Authors:** Samuele Di Carmine, Richard Cunningham, Katie J. Mylonas

## Abstract

Fibrosis develops in many organs through different initiating injuries, yet repeatedly engages inflammation, stromal activation and extracellular-matrix remodelling. Whether these recurring features represent a conserved disease programme, and whether commonly used mouse models reproduce it, remain unresolved. We developed ScarX, an R/Shiny workflow that analyses organ-resolved bulk RNA sequencing, identifies directional overlap across tissues, and ranks human-mouse conserved signals while retaining organ-restricted results. Public RNA-sequencing datasets representing fibrotic or fibrosis-associated disease in human heart, kidney, liver and lung were compared with matched mouse injury models. Differential expression was estimated independently within each organ using DESeq2, followed by cross-organ intersection, functional enrichment, matched-organ effect-profile comparison and a human-first conserved modulation (HCM) score that gives greatest weight to the breadth and magnitude of human disease change, adds support for same-direction mouse effects and penalises discordance.

In human disease, 202 genes increased and 35 decreased in the same direction across all four organs. Mouse models contained 409 pan-organ increases and 46 decreases. After gene-symbol harmonisation, 25 gene-direction signatures were conserved across all four organs in both species. These included *AEBP1*, *CDH11*, *COL1A1*, *COL1A2*, *COL3A1*, *COL5A2*, *COL6A2*, *EFEMP2*, *EMILIN1*, *LOXL1*, *LOXL2*, *TGFB2* and *VIM*. Matched human-mouse effect profiles varied considerably: Spearman rho was 0.42 for kidney, 0.31 for heart, 0.16 for liver and -0.06 for lung. Despite this uneven gene-level fidelity, 62 same-direction pathways were shared across species, led by extracellular-matrix organisation, collagen fibril organisation, cell-substrate adhesion, wound healing and ECM-receptor interaction. The HCM ranking concentrated structural matrix genes near the top but also identified cell-surface, secreted and regulatory candidates. Among the top 500 candidates, 376 carried at least one local tractability annotation.

ScarX therefore separates a compact conserved matrix-remodelling core from a much larger organ- and species-dependent response. The results provide a transparent starting point for target and biomarker selection, while also showing why expression conservation alone cannot establish therapeutic dependence.

## Introduction

Fibrosis arises when repair mechanisms persist after repeated injury, progressively replacing functional tissue with remodelled extracellular matrix. It is a major cause of chronic lung disease, cirrhosis, chronic kidney disease and heart failure[1–3]. Initiating lesions affect different cells and evolve over different timescales[1, 2, 4], but established disease repeatedly involves inflammatory-cell accumulation, fibroblast activation, altered parenchymal-stromal communication, matrix deposition and tissue stiffening[4, 5]. Whether these features form a common molecular programme remains central to the search for treatments with activity across fibrotic diseases.

Across organs, transforming growth factor beta (TGF-β) signalling, integrin-mediated adhesion, growth-factor responses, matrix cross-linking and mechanosensitive feedback recur[1, 3–5]. Collagen accumulation is also common, although the contributing genes and cell populations vary by tissue[1, 6–8]. Shared pathway enrichment therefore indicates convergence but does not establish that the same target is active or therapeutically relevant in every organ.

Cross-tissue single-cell studies have identified recurrent fibroblast lineages and activated states[9], including shared inflammation- and matrix-associated states alongside tissue-specific adaptations[10] and related stromal populations[11]. More than 12,000 human heart, lung, liver and kidney allograft samples also revealed pan-organ signatures of dysfunction and fibrosis[12]. These studies establish similarities across tissues but not a conserved programme in primary fibrotic disease.

Existing studies do not establish which genes change in the same direction across non-transplant fibrosis of the heart, kidney, liver and lung, or whether such changes are shared dependencies rather than consequences of mature scar. Fibroblast atlases focus on one lineage, whereas allograft studies include rejection, ischaemia-reperfusion injury and immunosuppression. Single-organ studies localise transcriptional programmes to scar-associated macrophages, mesenchymal cells, fibroblasts, immune cells and epithelial states[13–21], but do not establish which changes are conserved across all four organs.

Bulk transcriptomics captures the complete organ response across larger cohorts, but a signal can arise from altered expression within resident cells, expansion or recruitment of expressing cells, loss of other populations, or a combination of these processes[22, 23]. Bulk data alone cannot distinguish among these explanations or establish that increased matrix-gene expression represents deposited matrix or therapeutic dependence. LOXL2 illustrates the limitation: a randomised, double-blind, placebo-controlled phase 2 trial of simtuzumab (a LOXL2 inhibitor) in patients with idiopathic pulmonary fibrosis was stopped because it did not improve progression-free survival, including among participants with elevated serum LOXL2; the trial did not establish why inhibition failed[24]. Cross-organ analysis must therefore retain effect direction, magnitude and breadth and distinguish structural markers from candidates with plausible experimental or clinical use.

Experimental models add further uncertainty. Similarity between human chronic liver disease and mouse models depends on model and injury duration[25]. Spatial analysis distinguished arrested alveolar regeneration in idiopathic pulmonary fibrosis from active repair after bleomycin injury[26], and age alters whether bleomycin-induced fibrosis persists or resolves[27]. Because most comparisons cover one organ, it remains unclear whether mouse-human fidelity is organ dependent.

We therefore used ScarX to identify genes changing in the same direction across established human fibrosis of the heart, kidney, liver and lung and to test how closely matched mouse injury models reproduce them. We retained both the strict four-organ intersection and strong organ-restricted signals, assessed gene- and pathway-level overlap, quantified matched mouse-human effect-profile similarity and ranked conserved signals by magnitude, organ breadth and potential accessibility. The aim was to prioritise targets and biomarkers with potential relevance across organ-specific fibrotic diseases, rather than a single multisystem disorder. We hypothesised that a small, shared gene set would sit within broader organ- and species-specific change.

## Materials and Methods

### Study design

Public bulk RNA-sequencing datasets were assembled for disease-control comparisons in human heart, kidney, liver and lung and for matched injury-model comparisons in mouse. Human and mouse data followed the same count-based analytical workflow but were processed separately, with each organ analysed before cross-organ integration. The resulting organ-level summaries were linked through harmonised gene symbols, allowing directional conservation and effect-profile similarity to be assessed without pooling measurements generated in different species or studies.

This retrospective study used independently generated datasets to identify patterns for further testing. The source studies differed in design, disease stage, sampling and library preparation, making direct pooling of their raw samples inappropriate. Each dataset was analysed separately, and the resulting effect estimates were compared across organs. Disease-control contrasts were defined using the metadata accompanying each public study and retained in the ScarX input matrices.

### Public datasets

Public records were obtained from the Gene Expression Omnibus (GEO), Sequence Read Archive (SRA) and BioProject[28, 29]; the accessions represented in the current ScarX outputs are listed in Table 1. The GSE213001 lung series comprises 139 region-specific tissue records from 43 donors and includes idiopathic pulmonary fibrosis, non-diseased donor, non-IPF interstitial lung disease and chronic lung allograft dysfunction specimens[30]. ScarX used the pre-specified subset and condition labels in its bundled metadata. Several regions can originate from the same donor, so specimen numbers do not represent independent patient counts. The kidney comparison used advanced diabetic nephropathy and control tissue from GSE142025[31]. Cirrhotic and non-cirrhotic liver samples were drawn from GSE148355, a larger hepatocellular-carcinoma study[32], while the cardiac comparison paired ischaemic cardiomyopathy with non-failing left ventricle from GSE116250[33].

**Table 1.** Public RNA-sequencing datasets used in ScarX.

| Species / organ | Disease or model contrast | Public accession(s) | Records represented in the current analysis |
|---|---|---|---|
| Human / lung | Idiopathic pulmonary fibrosis-related lung tissue versus non-diseased donor lung | GSE213001;<br>SRP396424;<br>PRJNA878759 | Source series: 139 regional specimens from 43 donors; |
| Human / kidney | Advanced diabetic nephropathy versus control kidney | GSE142025;<br>SRP237545;<br>PRJNA595590 | 20 advanced diabetic nephropathy; 9 control |
| Human / liver | Cirrhotic versus normal liver tissue | GSE148355;<br>SRP255859;<br>PRJNA624047 | 9 cirrhotic; 13 normal |
| Human / heart | Ischaemic cardiomyopathy versus non-failing left ventricle | GSE116250;<br>SRP151309;<br>PRJNA477855 | 13 ischaemic cardiomyopathy; 12 non-failing |
| Mouse / lung | Bleomycin injury versus PBS control | SRP410457;<br>PRJNA906540;<br>GSM6762010-GSM6762023 | 5 bleomycin; 9 PBS |
| Mouse / kidney | Ischaemia-reperfusion injury versus sham | SRP504735;<br>PRJNA1105920 | 3 injury; 3 sham |
| Mouse / liver | Carbon tetrachloride injury versus oil control | SRP419015;<br>PRJNA927283 | 4 carbon tetrachloride; 4 oil |
| Mouse / heart | Coronary ligation/remodelling versus sham operation | SRP624724;<br>PRJNA1332716 | 3 ligation; 4 sham |

The mouse datasets represented four tissue-specific injury models. Lung samples compared bleomycin-treated mice with vehicle controls in GSE218997, a study of fibrosis resolution in young and old mice[27]. Kidney samples compared ischaemia–reperfusion injury with sham surgery in GSE266242, from a study of PFKFB3 in renal fibrosis[34]. Liver samples compared carbon tetrachloride with oil controls in GSE223686, a bulk RNA-sequencing series on thyroid hormone receptor β1 and fibrosis resolution. Heart samples compared myocardial infarction induced by coronary ligation with sham surgery in GSE308783[35]. For each tissue, ScarX analysed the pre-specified subset and condition labels in its bundled metadata; the sample counts in Table 1 refer to those selected records.

### Read acquisition, alignment and gene counting

Sample records in GEO were linked to their SRA run accessions and organised in species-specific manifests. Run archives were retrieved with SRA Toolkit prefetch, and fasterq-dump - -split-files generated FASTQ files while preserving paired-end structure. Read quality was reviewed from FastQC, MultiQC and, where available, optional fastp reports; these files were retained separately from the statistical outputs.

Rsubread was used to align reads against species-specific reference genomes[36]. Human libraries were mapped to the GRCh38 primary assembly with GENCODE release 44 annotation, whereas mouse libraries used GRCm39 and the NCBI RefSeq annotation for GCF_000001635.27. Reference indices were built once with Rsubread::buildindex. Where required, BAM files were sorted, indexed and summarised with samtools flagstat and samtools idxstats. Gene counts were obtained with Rsubread::featureCounts by assigning reads to exon features and aggregating them by gene_id[37]. Single-end and paired-end libraries were counted with the appropriate settings before count tables were merged over the union of gene identifiers. Missing gene-sample entries created during this within-dataset merge were treated as structural zeros, representing no reads assigned to that gene in that sample rather than model-inferred expression. These raw integer counts entered the organ-specific matrices; genes with fewer than 10 reads across the complete organ-specific comparison were removed before DESeq2 model fitting.

### Organ-wise differential expression

ScarX was implemented in R and deployed through Shiny. Within each organ, samples carrying the configured control and disease or injury labels were analysed with DESeq2[38]. Genes contributing fewer than 10 reads across the complete organ-specific comparison were removed before fitting the negative-binomial model with the design ∼ condition. Size-factor normalisation, dispersion estimation and model fitting followed the DESeq2 defaults. Log2 fold changes were then shrunken with apeglm, reducing unstable estimates for genes with limited information[39], and false-discovery rates were controlled with Benjamini-Hochberg adjusted P values.

Significance required both a Benjamini-Hochberg adjusted P value below 0.05 and an absolute shrunken log2 fold change above 0.20[40]. Positive and negative effects meeting these criteria were designated as upregulated and downregulated, respectively. ScarX retained the complete DESeq2 results alongside significant-gene tables and variance-stabilised expression values used for visualisation. The sample column supplied by each public study formed the primary unit of comparison. Some source series contain several regional specimens from one donor; because donor identity was not consistently available across studies, a donor-aware analysis remains a necessary sensitivity test.

### Cross-organ integration

Cross-organ integration began by joining the independently analysed organ results on annotated gene symbol. For each gene, ScarX recorded how many selected organs passed the significance threshold, how many effects were positive or negative, the mean and mean absolute log2 fold change across significant organs, and the smallest adjusted P value. A pan-organ increase required significant positive effects in heart, kidney, liver and lung; a pan-organ decrease required significant negative effects in all four. Records that did not meet either four-organ definition remained available as partially shared, mixed-direction or organ-restricted signals.

Venn and UpSet plots summarised the resulting overlaps, while the directionality matrix grouped genes by the number of organs with significant positive and negative effects.

### Functional enrichment

Functional enrichment was examined separately for positive and negative gene sets using Gene Ontology Biological Process and KEGG over-representation analysis in clusterProfiler[41]. Human and mouse symbols were mapped to Entrez identifiers through org.Hs.eg.db and org.Mm.eg.db. Terms were retained when the nominal P value was below 0.05 and the Benjamini-Hochberg q value was below 0.20. In the enrichment plots, dot sizeshows the number of genes assigned to a term, the horizontal position gives the gene ratio and colour represents the adjusted P value.

### Human-mouse conservation and model fidelity

Human and mouse organ summaries were aligned using gene symbols. For each gene, ScarX first calculated a human DiseaseScore and a mouse ModelScore that combined the number of organs meeting the significance criteria with effect magnitude and statistical support:

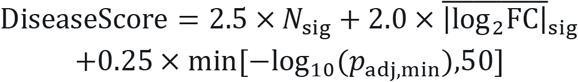

The human DiseaseScore and mouse ModelScore were calculated identically: 2.5 times the number of tissues with significant differential expression (Nsig), plus 2.0 times the mean of the absolute apeglm-shrunken log2 fold changes across those tissues, plus 0.25 times min[- log10(padj,min), 50]. Thus, the effect-size term is the mean of absolute effects rather than the absolute value of a signed mean. This effect-size term was not additionally capped; only the statistical-support term was capped at 50.

Matched genes were subsequently classified as conserved up, conserved down, discordant, human only, mouse only or mixed/weak. Conserved genes passed the active thresholds in both species and changed in the same direction; discordant genes were significant in both but changed in opposite directions. Human-only and mouse-only labels were assigned when the threshold was met in one species, and all remaining records were retained in the mixed/weak class.

The human-first conserved modulation (HCM) score then combined the human DiseaseScore and mouse ModelScore with the direction of cross-species agreement and the breadth of significant change:

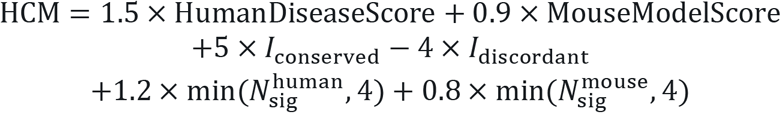

Cross-species direction was represented by binary terms. The conservation indicator was one when a gene changed in the same direction in human disease and the corresponding mouse model, whereas the discordance indicator was one for opposing changes; otherwise the relevant indicator was zero. The human DiseaseScore and human organ-breadth term were weighted more strongly than their mouse equivalents so that a strong model effect could not outweigh limited human support. The coefficients were prespecified for ranking and were not derived from experimental outcomes; HCM is therefore a prioritisation measure, not an estimate of effect size or therapeutic success.

Mouse-model fidelity was assessed from Spearman correlations between gene-level effects in each human and mouse organ pairing. Each matrix cell reports both the correlation coefficient and the number of matched genes included in the calculation. Genome-wide correlation was complemented by a focused analysis of genes reaching significance in at least one species within each matched organ. These genes were grouped as conserved up, conserved down, discordant, human only or mouse only, allowing broad transcriptomic similarity to be considered alongside agreement among the disease-associated signals.

### Shared-pathway analysis and candidate ranking

Shared functional programmes were identified by matching the human and mouse enrichment tables on database, term and direction. A pathway qualified as shared when it was significant in both species and carried the same direction label; qualifying terms were ordered using their false-discovery values in the two analyses.

Candidate plots placed the HCM ranking alongside conservation class, organ breadth and mean effect size. Experimental accessibility was explored by annotating genes encoding secreted proteins, cell-surface proteins, kinases and transcription factors. A gene could carry several annotations. For visualisation, overlapping annotations were collapsed into a single display class, while genes without an assigned class were retained. These categories identify candidates that may be accessible to measurement or perturbation.

### Target Board and provisional biomarker panel

The Target Board ranks each candidate according to the number of affected organs, the magnitude and statistical strength of its expression changes, consistency of direction and potential tractability. These features are combined in the following raw impact score:

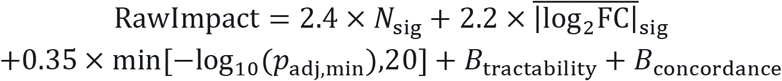

Variables shared with the DiseaseScore and HCM equations retain the definitions given above. The tractability bonus assigns 1.2, 1.1, 0.9 and 0.7 points to secreted, cell-surface, kinase and transcription-factor annotations, respectively. Fully concordant genes receive a further 2.5 points. For genes showing mixed directions across organs, the concordance bonus is the absolute difference between the numbers of upregulated and downregulated organs divided by the total number of significant organs. Cross-species analyses use HCM in place of this ranking. To aid comparison within the displayed candidate set, scores are expressed as a percentage of the highest value in the current view.

The provisional biomarker shortlist was based on organ breadth, effect magnitude, statistical support, directional consistency and annotation as a secreted or cell-surface protein. Selection used tissue and blood expression data available from the Human Protein Atlas[42]. Direct measurements are required to determine whether these proteins enter the circulation, can be detected across a suitable concentration range and distinguish fibrosis from other forms of tissue injury.

### Software implementation and reproducibility

Differential expression was performed with DESeq2, with apeglm used to shrink log2 fold-change estimates. Gene identifiers were mapped through AnnotationDbi and the corresponding human and mouse annotation databases, while functional enrichment was carried out with clusterProfiler. Figures were generated using ggplot2, pheatmap and EnhancedVolcano. ScarX also includes optional modules that access decoupleR, OmniPath and CollecTRI for transcription-factor activity, together with Ensembl, Open Targets and ChEMBL for therapeutic annotation (Badia-I-Mompel et al., 2022; Turei et al., 2021; Muller-Dott et al., 2023). These modules did not contribute to the bulk differential-expression results reported in this study.

## Results

### Human fibrotic tissues share a small directional core within a much broader disease response

All four human comparisons contained extensive differential expression at adjusted P below 0.05 and absolute shrunken log2 fold change above 0.20. Kidney had the largest set, with 8,868 increased and 7,241 decreased genes. Heart contained 5,250 increased and 4,587 decreased genes; lung contained 5,144 increased and 4,182 decreased genes; and liver contained 4,638 increased and 4,332 decreased genes (**Figure 1A**). Positive and negative changes were evident in every organ-level volcano plot. Kidney produced the broadest significant response, while heart, liver and lung retained distinct distributions of effect magnitude and statistical support (**Supplementary Figure S1A**). Correlations among the complete human organ profiles were modest, with lung showing the weakest overall similarity to the remaining tissues (**Supplementary Figure S1B**).

**Figure 1.**
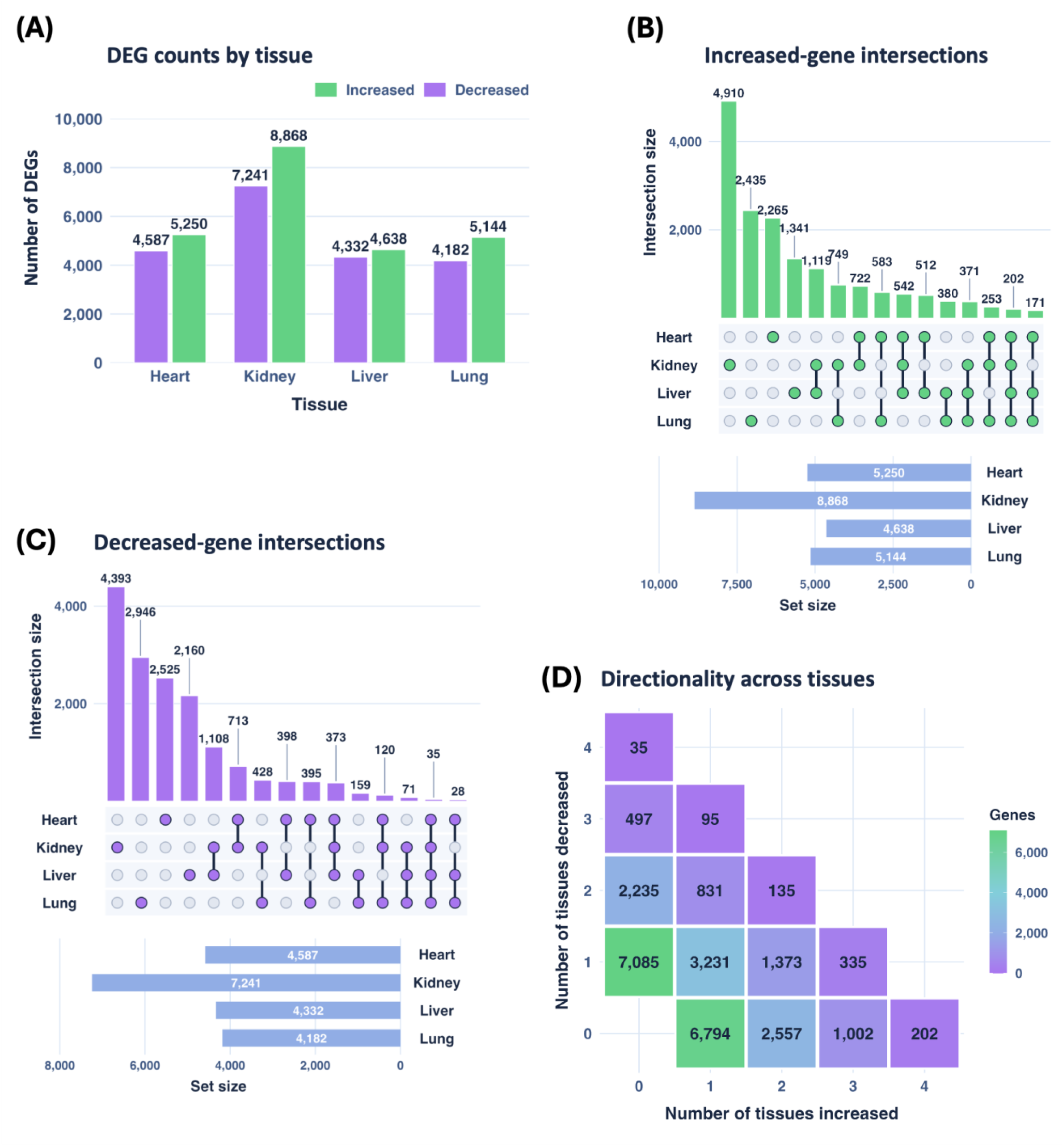
Human differential-expression burden and cross-organ overlap. **(A)** Numbers of significant increased and decreased genes in heart, kidney, liver and lung. **(B, C)** UpSet plots show intersection sizes for increased and decreased genes, respectively. **(D)** Each cell reports the number of genes assigned to a combination of organs with significant positive and negative change. Significance required adjusted P < 0.05 and shrunken log2 fold change > 0.20.

Cross-organ intersection reduced this broad response sharply. The increased-gene UpSet plot identified 202 genes shared by all four organs and much larger organ-specific sets, particularly in kidney (**Figure 1B**). The decreased-gene UpSet plot identified 35 genes shared by all four organs and showed a similar predominance of organ-specific change (**Figure 1C**). The directionality matrix contained 6,000 genes with mixed significant directions, 10,353 with one or more positive changes and no significant negative change, and 9,817 with one or more negative changes and no significant positive change (**Figure 1D**). The Venn diagrams confirmed that these four-organ intersections were small relative to the organ-restricted compartments. Kidney alone contained 4,910 increased and 4,393 decreased genes, compared with 202 increased and 35 decreased genes at the centre of the four-organ overlaps (**Supplementary Figure S2A**; **Supplementary Figure S2B**).

Among the ten human genes selected by significant-organ breadth and effect magnitude, the bold radar profile reports the organ-wise median signed apeglm-shrunken log2 fold change across the displayed genes; individual profiles show gene-level variation around it (**Supplementary Figure S3A**). Hierarchical clustering paired heart and kidney most closely, with liver joining next and lung forming the most distinct branch (**Supplementary Figure S3B**). The direction-flow plot showed extensive movement among upregulated, downregulated and non-significant classes across organs, with a much smaller fraction retaining the same state throughout the displayed sequence (**Supplementary Figure S3C**).

Taken together, the human comparisons contain a reproducible but compact pan-organ response within a much larger organ-dependent transcriptome. Shared increases were more common than shared decreases, while most genes and whole-profile relationships retained a strong tissue-specific component.

### The human pan-organ increase is dominated by matrix organisation and tissue remodelling

GO Biological Process enrichment of the 202 pan-organ increased genes was led by extracellular matrix organisation, extracellular structure organisation and external encapsulating structure organisation (**Figure 2A**). Collagen fibril organisation, collagen metabolic and biosynthetic processes, wound healing, cell-substrate adhesion and Wnt-related terms were also represented. The 35 pan-organ decreased genes produced a narrower GO set centred on oxidative phosphorylation, aerobic respiration, ATP metabolic processes and mitochondrial electron transport (**Figure 2B**).

**Figure 2.**
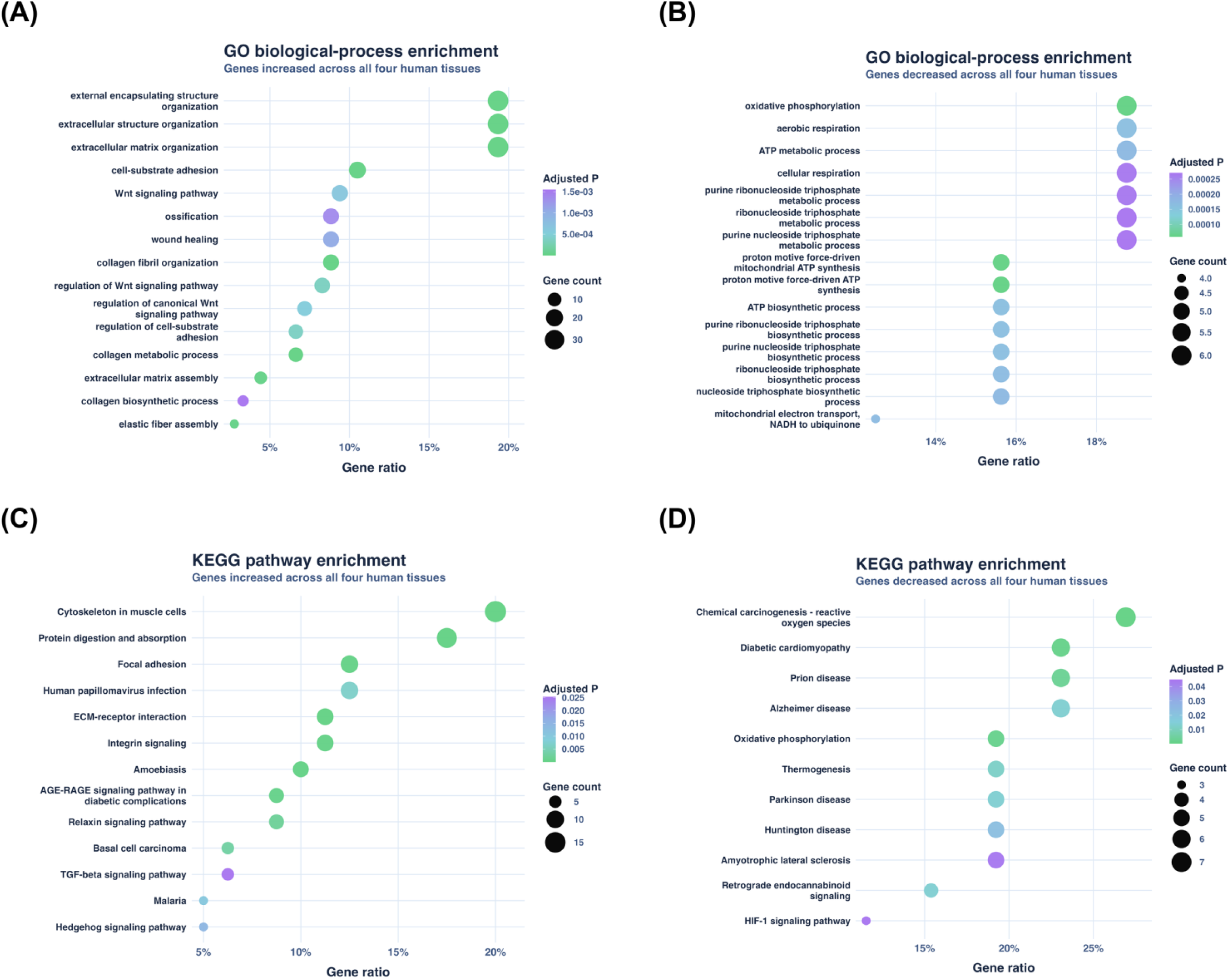
Functional enrichment of the human four-organ directional signatures. GO Biological Process **(A, B)** and KEGG **(C, D)** dot plots summarise terms enriched among genes increased in all four organs **(A, C)** and decreased in all four organs **(B, D)**. The x-axis gives the gene ratio, point size gives the contributing gene count and colour gives the adjusted P value.

KEGG enrichment of the 202 increased genes included cytoskeleton in muscle cells, protein digestion and absorption, focal adhesion, ECM-receptor interaction, integrin signalling, AGE-RAGE signalling and TGF-β signalling (**Figure 2C**). The decreased-gene KEGG terms similarly included oxidative phosphorylation and several neurodegenerative-disease pathways whose shared membership is driven by mitochondrial respiratory-chain genes (**Figure 2D**).

Together, the human four-organ signature combined a broad matrix-remodelling increase with a smaller reduction in oxidative metabolism. The imbalance in the size and breadth of these two signatures identifies extracellular-matrix organisation as the dominant shared feature of the established human disease tissues.

### Mouse injury models reproduce broad fibrotic features but retain model-specific transcriptional structure

The four mouse models showed extensive differential expression. Kidney ischaemia-reperfusion injury produced 5,135 increased and 4,870 decreased genes. Bleomycin lung injury produced 3,512 increased and 3,484 decreased genes; carbon tetrachloride liver injury produced 3,179 increased and 2,879 decreased genes; and post-ligation cardiac remodelling produced 3,009 increased and 1,651 decreased genes (**Figure 3A**). The volcano plots confirmed broad, bidirectional changes in every model, with kidney producing the largest and most statistically prominent response (**Supplementary Figure S1C**). Correlations between complete organ-level effect profiles were positive but modest, indicating shared and organ-specific components (**Supplementary Figure S1D**).

**Figure 3.**
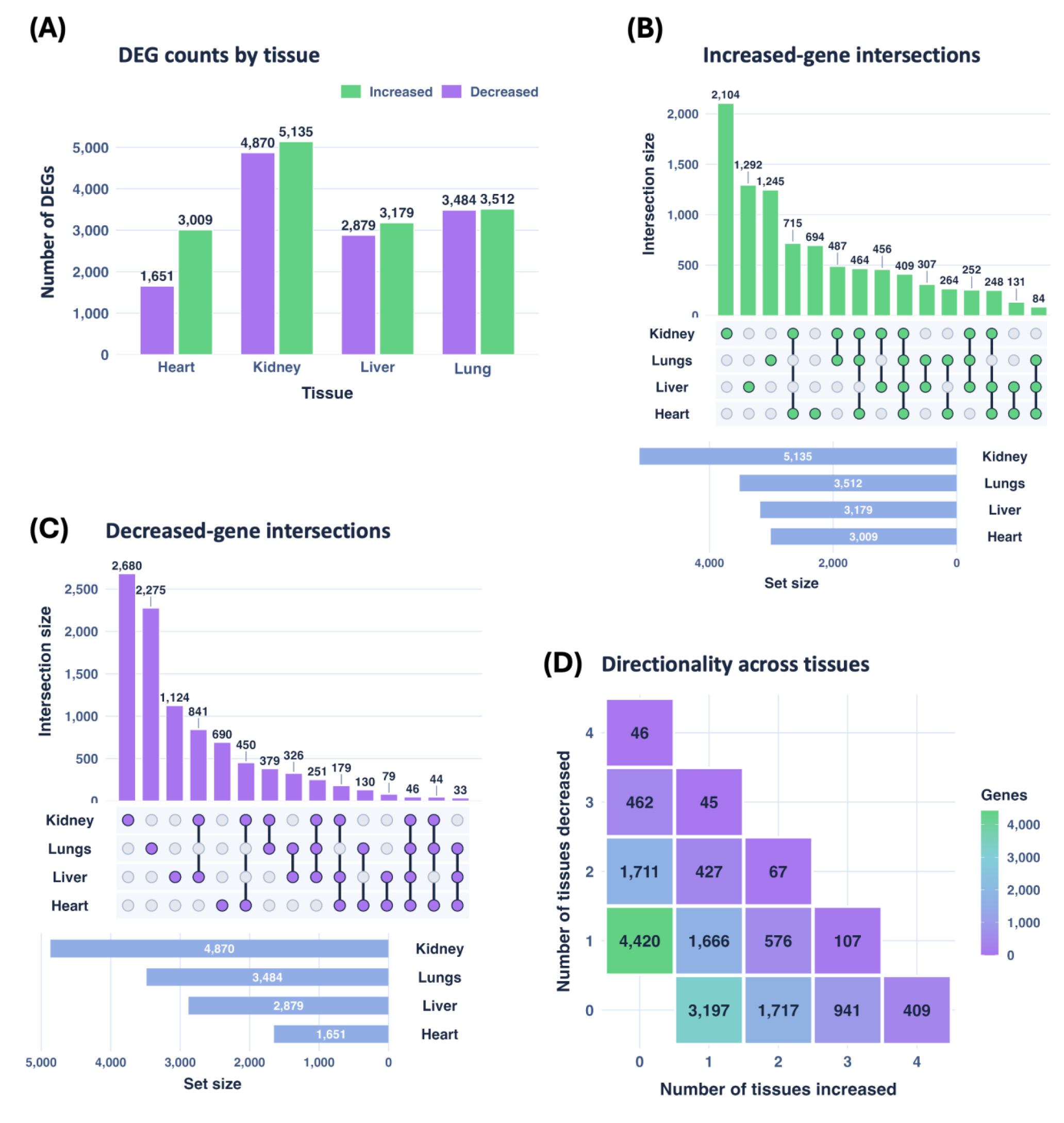
Mouse differential-expression burden and cross-organ overlap. **(A)** Numbers of significant increased and decreased genes in heart, kidney, liver and lung models. **(B, C)** UpSet plots show intersection sizes for increased and decreased genes. **(D)** Directionality matrix reports gene counts for each combination of significant positive and negative organ effects. Significance required adjusted P < 0.05 and shrunken log2 fold change > 0.20.

A total of 409 genes were increased in all four mouse organs (**Figure 3B**), whereas 46 were decreased across all four models (**Figure 3C**). The positive core was approximately twice the size of its human counterpart, while the negative core remained small in both species. Most significant genes were restricted to one organ or shared by only a subset. The directionality matrix contained 2,888 genes with mixed effects, 5,855 with significant increases and no decreases, and 6,593 with significant decreases and no increases (**Figure 3D**). The Venn diagrams placed the 409 increased and 46 decreased genes within much larger partial and organ-restricted intersections. Kidney contributed the largest organ-specific set in both directions, while lung also contained a substantial organ-restricted decrease (**Supplementary Figure S4A**; **Supplementary Figure S4B**).

The ten highest-ranked mouse genes generally retained broad effects across the four models. The bold radar profile reports the organ-wise median signed apeglm-shrunken log2 fold change across the displayed genes, with individual profiles showing gene-level variation (**Supplementary Figure S5A**). Correlation-based clustering placed kidney and liver closest together, with lung joining before heart, which had the most distinct profile (**Supplementary Figure S5B**). The direction-flow plot showed frequent movement among upregulated, downregulated and non-significant classes, with fewer genes retaining one classification throughout the displayed organ sequence (**Supplementary Figure S5C**).

Together, these results identify a measurable pan-organ response in the mouse models, dominated by increased rather than decreased expression. This shared core sits within a much larger model-dependent response: most genes were organ restricted, cross-organ correlations remained modest and direction frequently changed between tissues. The four models therefore reproduce common components of fibrotic injury while preserving the distinct transcriptional context of each organ.

### Mouse pan-organ enrichment combines inflammatory recruitment with matrix remodelling

The mouse pan-organ positive set was enriched for leukocyte migration, regulation of innate immune responses, myeloid leukocyte activation, phagocytosis, wound healing and extracellular-matrix organisation (**Figure 4A**). The mouse pan-organ negative set was enriched for carboxylic-acid biosynthesis, organic-acid biosynthesis, amino-acid metabolism, sulphur-compound biosynthesis and related metabolic processes (**Figure 4B**). These terms contrasted with the inflammatory, phagocytic and matrix-associated functions enriched in the positive set.

**Figure 4.**
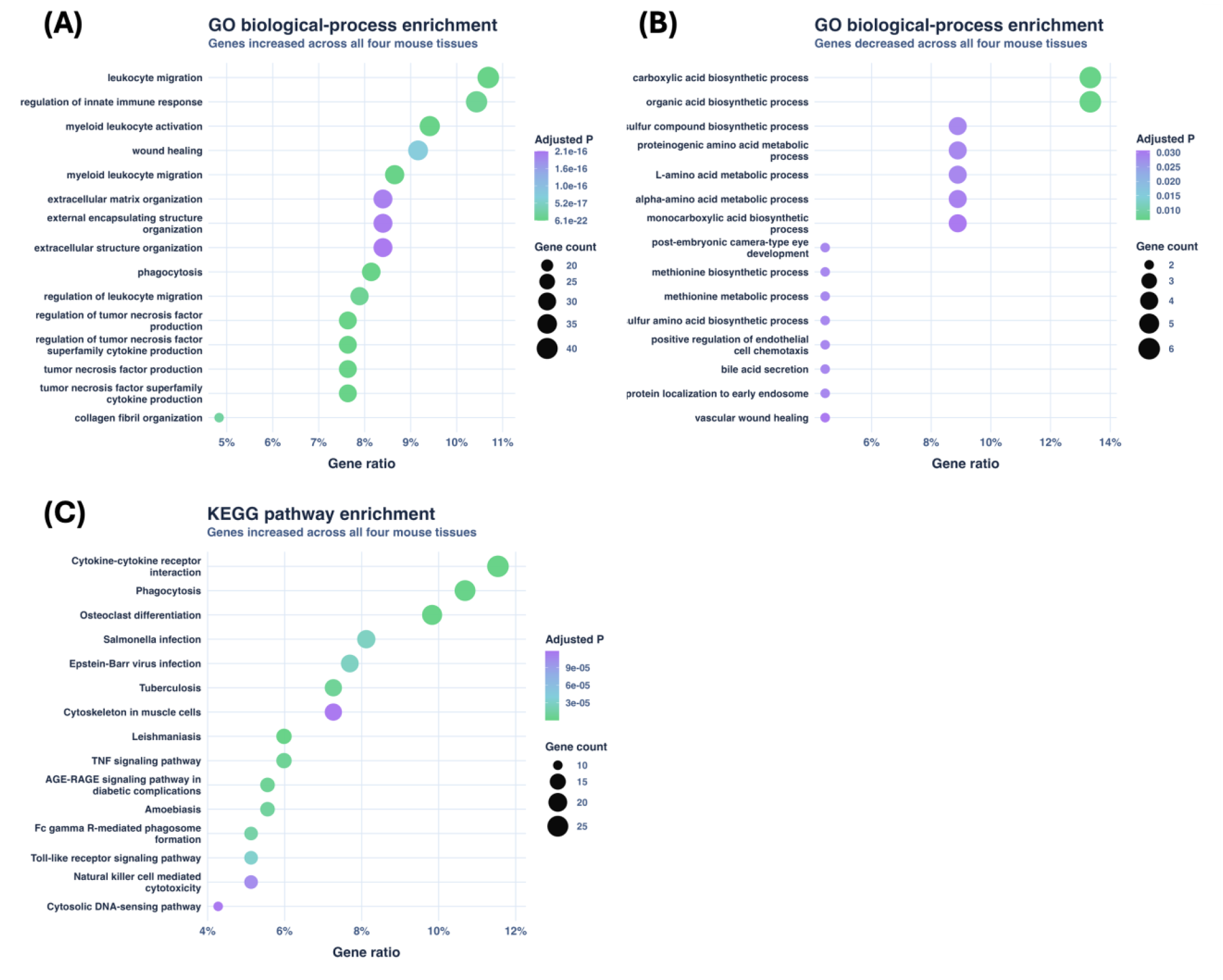
Functional enrichment of the mouse four-organ directional signatures. GO Biological Process dot plots summarise terms enriched among genes increased **(A)** or decreased **(B)** in all four models. The KEGG plot **(C)** shows terms enriched among the pan-organ increased genes. The x-axis gives the gene ratio, point size gives the contributing gene count and colour gives the adjusted P value.

KEGG enrichment of the mouse pan-organ positive set included cytokine-cytokine receptor interaction, phagocytosis, osteoclast differentiation, TNF signalling, Toll-like receptor signalling, AGE-RAGE signalling and cytosolic DNA sensing (**Figure 4C**). Several infection-labelled pathways were also returned because their host-response gene sets contain cytokine, phagocytic and cytoskeletal components.

The shared mouse programme therefore couples inflammatory recruitment and matrix remodelling with reduced biosynthetic and metabolic activity. Its prominent innate-immune component differentiates it from the more strongly matrix-centred human positive signature.

### Cross-species conservation is strongest in kidney and heart and weakest in the lung comparison

Within matched organs, the proportion of significant signals assigned to conserved up or conserved down varied substantially (**Figure 5A**). Kidney had the largest conserved fraction: 1,969 conserved increases and 1,882 conserved decreases, together accounting for 35.2% of genes significant in at least one species. The corresponding conserved fractions were 17.0% in heart, 14.9% in liver and 13.8% in lung. Lung also had the largest discordant fraction, 15.9%, compared with 4.8% in heart, 8.9% in kidney and 7.1% in liver.

**Figure 5.**
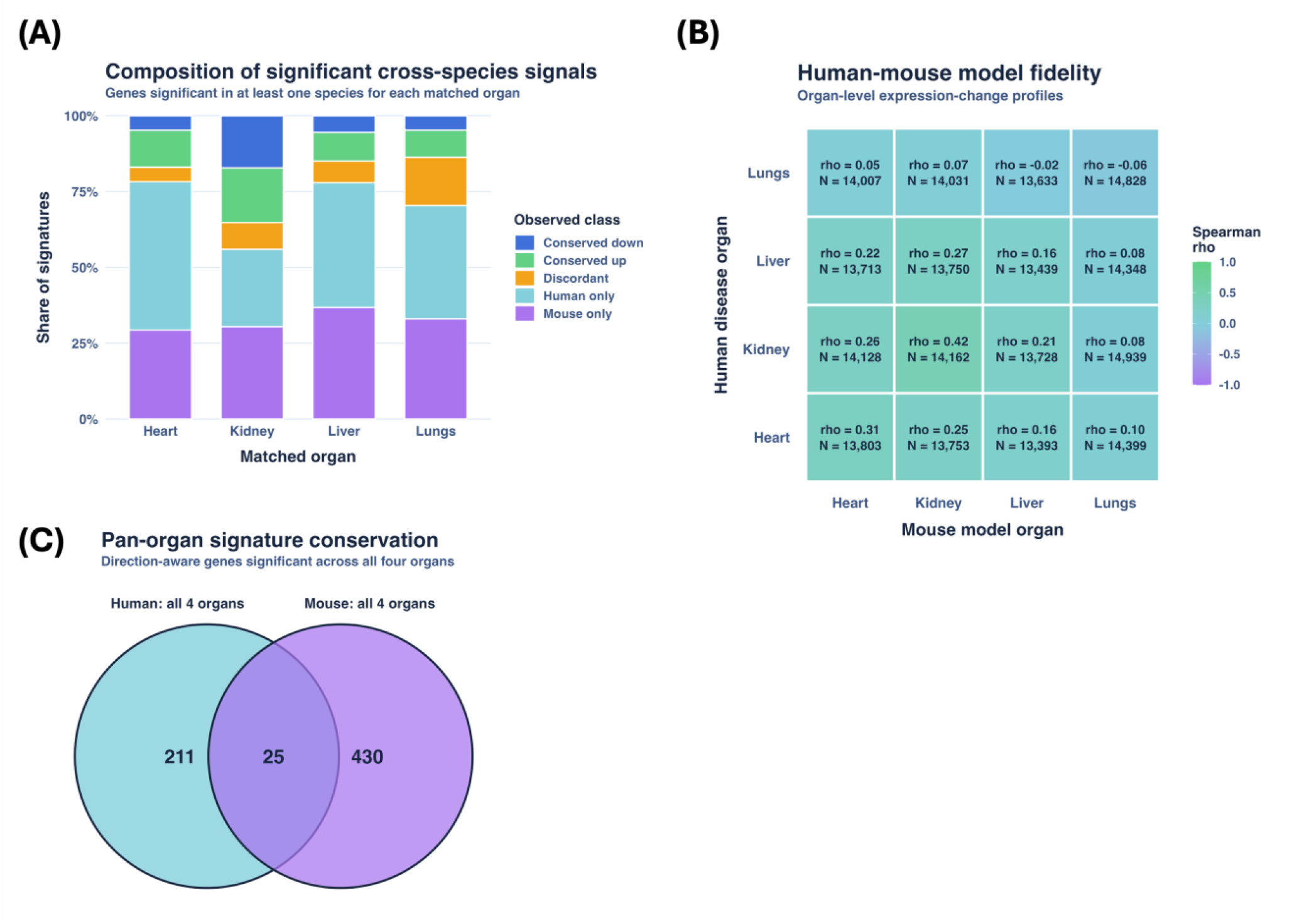
Human-mouse conservation across matched organs. **(A)** Percentage composition of conserved-down, conserved-up, discordant, human-only and mouse-only signals among genes significant in at least one species for each matched organ. **(B)** Spearman correlations between complete matched gene-effect profiles for every human-organ by mouse-organ pair; N is the number of matched genes. **(C)** Overlap of pan-organ gene-direction records after gene-symbol harmonisation.

Effect-profile correlations across all matched genes supported the same uneven pattern (**Figure 5B**). The matched kidney comparison was highest (rho=0.422; 14,162 genes), followed by heart (rho=0.310; 13,803 genes) and liver (rho=0.163; 13,439 genes). The matched lung value was slightly negative (rho=-0.062; 14,828 genes). Some off-diagonal correlations equaled or exceeded the matched liver and lung values, indicating that organ identity alone did not guarantee the closest transcriptome-wide match.

After gene-symbol harmonisation, the human data contained 236 pan-organ gene-direction records and the mouse data contained 455. Twenty-five were present in both species with the same direction across all four organs (**Figure 5C**; **Supplementary Table S1**). These were *AEBP1*, *B3GNT9*, *CACNB3*, *CDH11*, *COL1A1*, *COL1A2*, *COL3A1*, *COL5A2*, *COL6A2*, *DBN1*, *EFEMP2*, *EMILIN1*, *FBLN2*, *FKBP10*, *FKBP4*, *GPR176*, *KDELR3*, *LOXL1*, *LOXL2*, *OLFML2B*, *PMEPA1*, *RAB7B*, *SPECC1*, *TGFB2* and *VIM*. The overlap was enriched for matrix, matrix-processing and stromal-associated genes, with additional membrane, trafficking and signalling candidates. The four-set Venn diagram placed these signatures within much larger matched-organ sets. Kidney had the largest organ-restricted same-direction group, with 2,812 genes, compared with 744 in lung, 624 in heart and 574 in liver (**Supplementary Figure S6A**).

Cross-species conservation was strongest in kidney and heart, yet no organ pair fully reproduced the corresponding human disease profile. The 25-signature intersection defines a compact shared group surrounded by substantially larger organ- and model-specific responses.

### Conserved pathway structure is broader than the exact pan-organ gene intersection

Sixty-two enriched terms were significant in human and mouse and moved in the same direction. The strongest shared terms were extracellular matrix organisation, external encapsulating structure organisation, extracellular structure organisation and collagen fibril organisation (**Figure 6A**). Cell-substrate adhesion, extracellular-matrix assembly, elastic-fibre assembly, integrin signalling, wound healing, collagen biosynthesis and ECM-receptor interaction were also shared. The broader pathway overlap relative to the 25-gene exact intersection reflects partially different genes contributing to related processes in each species.

**Figure 6.**
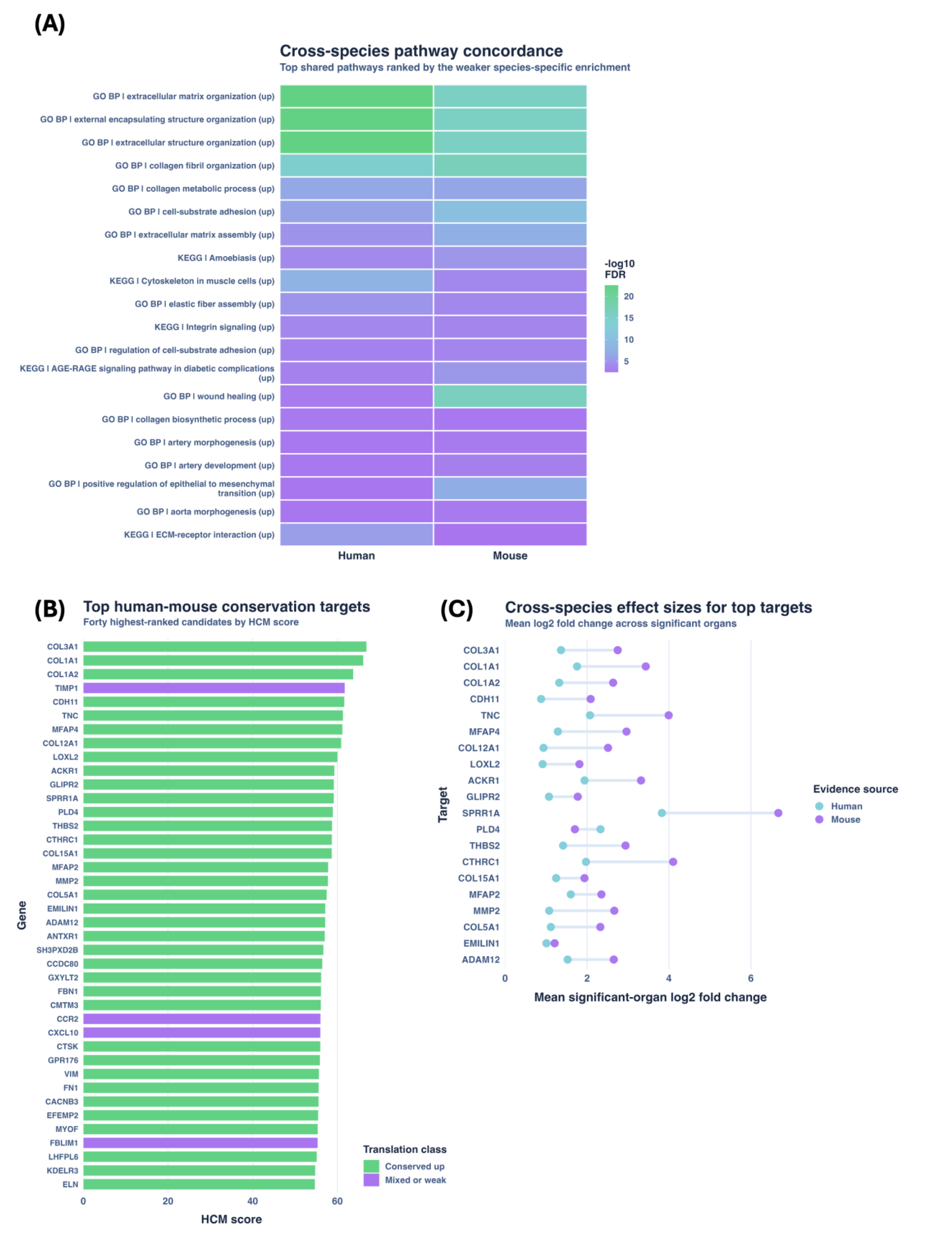
Shared pathways and HCM-ranked candidates. **(A)** Same-direction GO Biological Process and KEGG terms significant in both species; fill is -log10 false-discovery rate. **(B)** Leading HCM-ranked genes coloured by conservation class. **(C)** Human and mouse mean log2 fold changes among significant organs for selected high-ranking targets; the connecting line displays the species difference.

HCM ranking placed *COL3A1*, *COL1A1* and *COL1A2* first, with scores of 66.90, 66.14 and 63.76 (**Figure 6B**). *CDH11*, *TNC*, *MFAP4*, *COL12A1* and *LOXL2* also ranked highly and were classified as conserved increases. TIMP1 ranked fourth because of its broad, large signal but remained labelled mixed/weak under the conservation rules.

For the leading candidates, mouse mean effects were often larger than human effects (**Figure 6C**). *COL1A1*, *COL3A1*, *TNC*, *CTHRC1* and several other matrix-associated genes showed this translation gap. The dumbbell display retains the two species estimates, allowing a high HCM score to be examined alongside its effect-size imbalance.

These analyses place the strongest cross-species agreement at the level of matrix-related pathways, while individual genes retain differences in conservation class and effect magnitude. A high HCM rank therefore identifies a strong comparative signal whose human and mouse components still need to be considered separately.

### Conservation breadth and tractability annotations separate markers from possible intervention points

Across all matched genes, same-direction conservation was most often observed in one, two or three organs, while only 25 genes were conserved across all four (**Figure 7A**). These 25 genes formed a small vertical stratum within the broader conservation landscape. Structural collagens occupied the highest HCM range, whereas *CDH11*, *LOXL2*, *TGFB2*, *ACKR1*, *GPR176* and other non-collagen candidates combined cross-species conservation with secreted, cell-surface or other tractability annotations.

**Figure 7.**
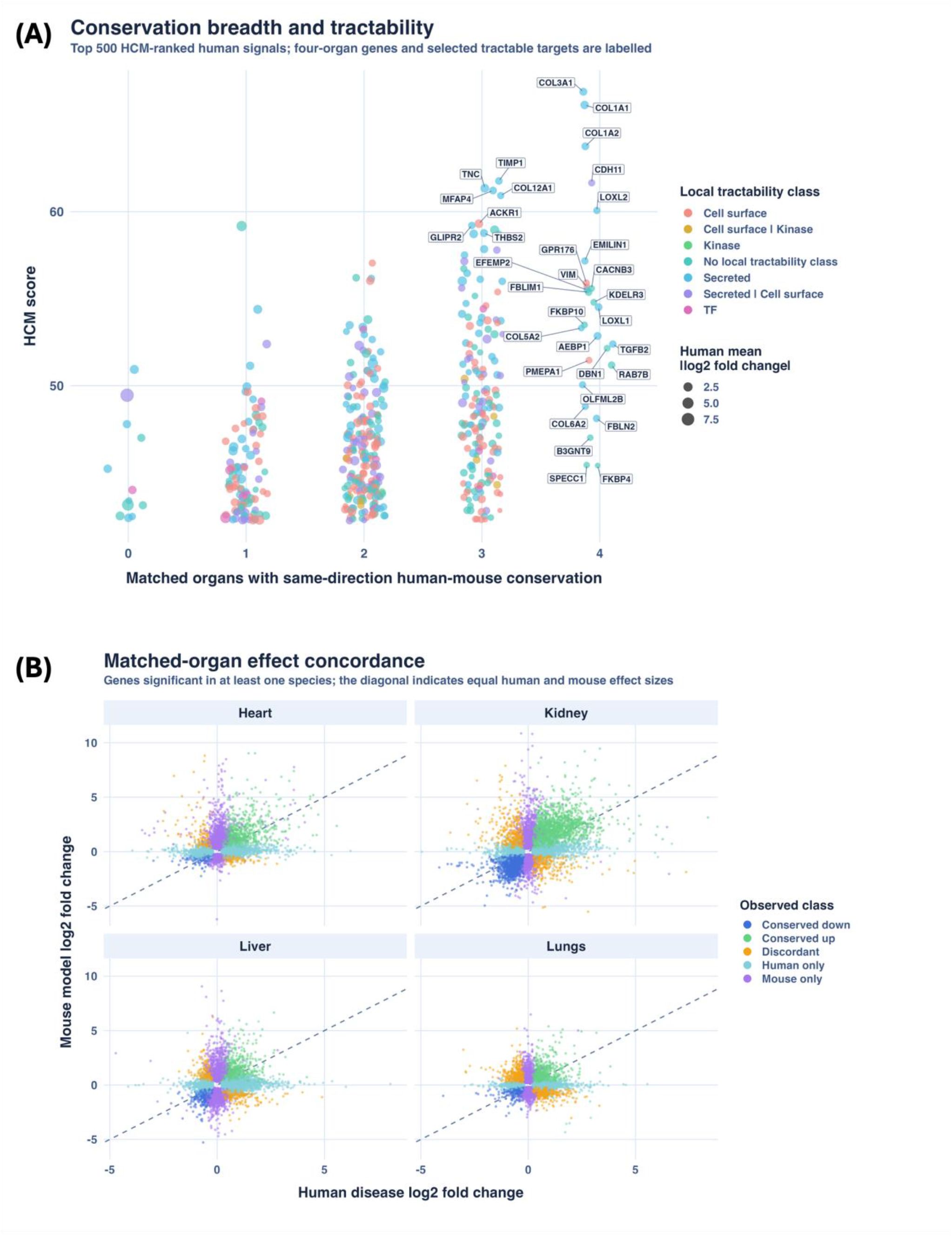
Conservation breadth, effect size and local tractability annotation. **(A)** HCM score plotted against the number of matched organs with same-direction human-mouse conservation. Colour indicates the local tractability class and point size indicates human mean absolute log2 fold change. **(B)** Human and mouse log2 fold changes for genes significant in at least one species, separated by matched organ and coloured by translation class; the diagonal marks equal effects.

The strict four-organ intersection represents the most consistently conserved signals, but it excludes candidates with strong human disease effects and conservation in fewer organs. Tractability was therefore examined across the 500 highest HCM-ranked genes, including the four-organ core and candidates with partial cross-species conservation. Of these, 376 (75.2%) had at least one displayed accessibility annotation.

The matched-organ scatter plots compared human and mouse effect sizes independently of conservation breadth (**Figure 7B**). Conserved positive genes occupied the positive-positive region, alongside substantial human-only, mouse-only and discordant populations. Kidney contained the broadest same-direction distribution, whereas lung showed a larger discordant component. In the aggregate analysis, most genes had modest effects in both species. Conserved increases extended towards larger mouse effects, while mixed, weak and discordant genes remained concentrated at the lower effect range. Larger HCM points occurred mainly among genes with stronger effects and conservation across several organs (**Supplementary Figure S6B**).

Conservation breadth, effect size and tractability therefore captured distinct features of the ranked candidates. Structural genes accounted for many of the most recurrent signals, while several non-collagen candidates combined cross-species agreement with annotations that may support experimental testing.

### Target and biomarker views expose the distinction between ranking and validation

The relative estimated-impact display in the Target Board ranked *COL3A1* first, with *COL1A1*, *COL1A2*, *CDH11* and *LOXL2* also among the leading candidates (**Figure 8A**). The percentage is normalised to the top candidate in the displayed set and therefore indicates relative priority inside the current ScarX view. It does not represent the probability or expected magnitude of therapeutic benefit.

**Figure 8.**
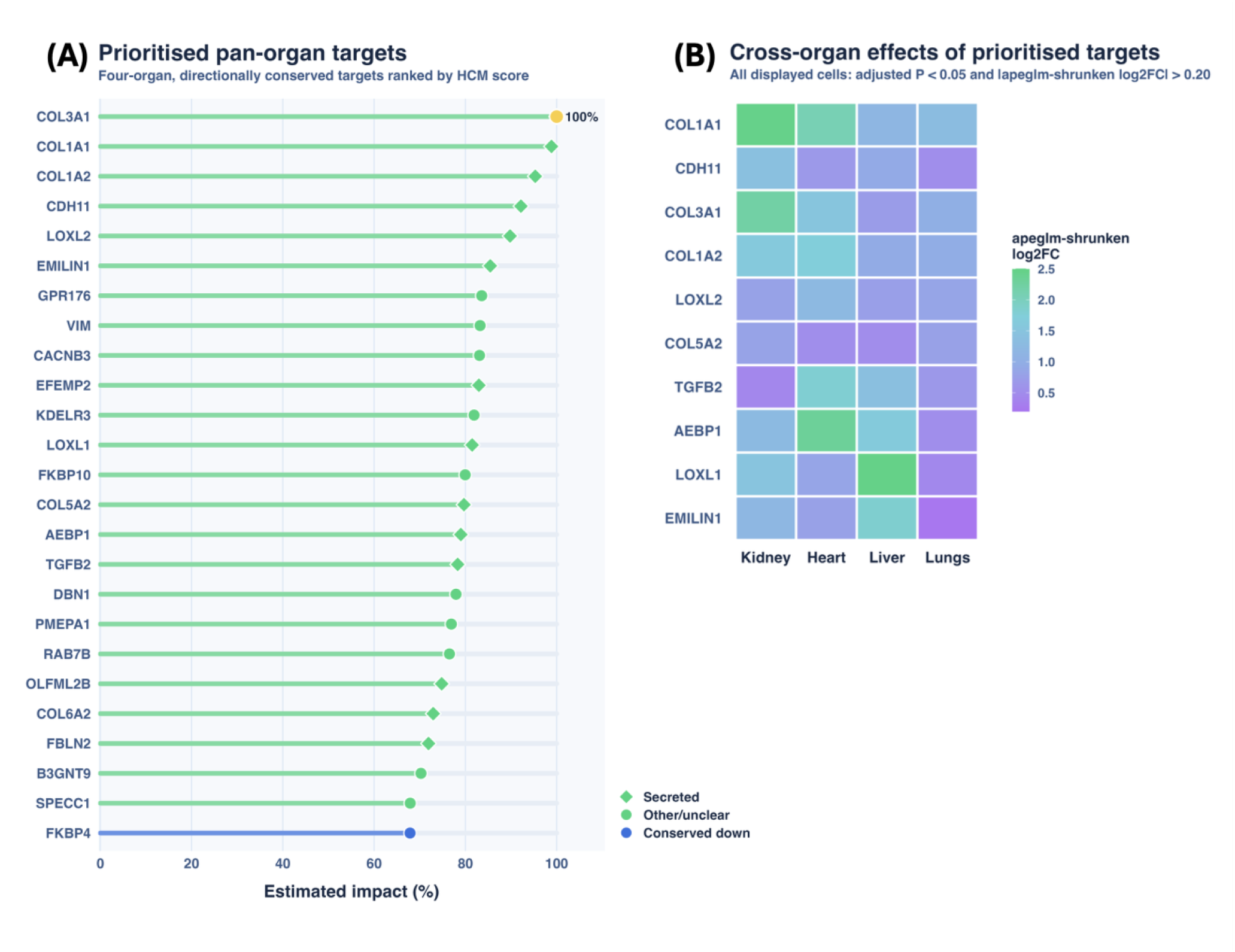
ScarX target and provisional biomarker views. **(A)** Relative estimated-impact scores for the displayed Target Board candidates, normalised to the highest score in the view; symbols report available annotations. **(B)** Organ-resolved human apeglm-shrunken log2 fold changes for a provisional ten-gene panel; all displayed gene-organ combinations met the ScarX significance threshold (adjusted P < 0.05 and absolute shrunken log2 fold change > 0.20).

A ten-gene provisional panel was visualised across the four human organs using apeglm-shrunken log2 fold changes (**Figure 8B**). The panel contained *COL1A1*, *CDH11*, *COL3A1*, *COL1A2*, *LOXL2*, *COL5A2*, *TGFB2*, *AEBP1*, *LOXL1* and *EMILIN1*. All 40 displayed gene-organ combinations met both ScarX significance criteria (adjusted P < 0.05 and absolute shrunken log2 fold change > 0.20), so the redundant cell outlines were removed. All ten candidates increased across kidney, heart, liver and lung, although the magnitude varied. The Human Protein Atlas reports each encoded protein as detectable in blood, providing a basis for considering a provisional circulating panel.

In conclusion, the Target Board identifies conserved genes for functional investigation, while the biomarker panel extends the analysis towards blood-based measurement. Together, they define a focused group of candidates for experimental perturbation and biomarker validation.

## Discussion

Using ScarX, we defined a compact transcriptional programme shared across fibrotic or fibrosis-associated disease in human heart, kidney, liver and lung and the corresponding mouse injury models. The human datasets contained 202 genes increased across all four organs and 35 decreased, compared with 409 increases and 46 decreases in mice. Requiring the same directional change in every organ and both species narrowed this overlap to 25 gene signatures, while pathway analysis identified 62 shared terms centred on adhesion, wound repair and matrix-receptor signalling. Complete gene-effect profiles were less concordant: kidney and heart showed the strongest matched-organ correlations, liver was weaker and lung was slightly negative. These findings support a shared matrix-centred core within a much larger organ-, disease- and model-dependent response.

Extracellular-matrix genes were prominent within the core, consistent with recurrent collagen deposition, myofibroblast activation, matrix organisation and mechanical signalling across fibrotic diseases[1–3]. TGF-β, integrin, Wnt, growth-factor and matrix-remodelling pathways also recur after different forms of tissue injury[4, 5]. Yet only 25 genes retained direction across all eight comparisons despite thousands of changes within individual tissues. The shared response is therefore real but forms a small component of the broader organ-specific transcriptome.

Pathway-level agreement was broader than gene-level agreement because related processes can recur through different genes or cell populations. The 62 terms repeatedly included matrix organisation, cell adhesion, collagen assembly and wound healing. Cross-tissue single-cell studies have similarly identified related fibroblast and fibro-inflammatory states without complete agreement among marker genes[9–11, 13, 14, 16–19]. Pathway convergence therefore complements the strict gene intersection but does not establish that every organ uses the same molecular route.

Model fidelity also varied. Kidney showed the closest overall correspondence and largest conserved fraction, heart retained moderate concordance, liver was weaker and lung showed little whole-profile similarity despite a subset of conserved matrix genes. Bleomycin injury can reproduce active fibrotic signals while undergoing resolution that distinguishes it from progressive human disease[27, 43], and liver-model conservation varies by pathway, stage and experimental context[25]. Sampling region, disease duration, age, cellular composition and technical design may also influence the correlations. Candidate relevance must therefore be interpreted alongside the broader fidelity of the model used for testing.

The HCM score was designed for that purpose. It gives greatest weight to the breadth and magnitude of human disease effects, adds support for same-direction mouse effects and penalises discordance. Each rank must be read alongside the underlying organ-level effects, conservation class, biological function and tractability.

Structural genes such as *COL3A1*, *COL1A1* and *COL1A2* ranked highly because their changes were large, recurrent and directionally conserved. Their abundance makes them strong markers of matrix accumulation but does not establish a safe or effective therapeutic target. The clinical trial cited above illustrates that expression conservation alone does not establish therapeutic dependence[24]. HCM should therefore be used to prioritise experiments, not predict therapeutic success.

Tractability annotations extended the candidate set beyond fibrillar collagens: 376 of the top 500 HCM-ranked genes encoded a secreted, cell-surface, kinase or transcriptional-regulatory protein. CDH11 already has direct intervention evidence: genetic deletion and a neutralising antibody reduced bleomycin-induced pulmonary fibrosis, and CDH11-mediated contact between macrophages and myofibroblasts localised latent TGF-β activation[44, 45]. Such annotations guide assay design but are not equivalent to druggability; localisation, isoform specificity, tissue distribution, safety, compound quality and assay context remain essential.

*TGFB2* connects the ranking to an established profibrotic pathway. An active enhancer sustains autocrine TGF-β2 signalling in systemic sclerosis fibroblasts; NF-κB or BRD4 inhibition reduced profibrotic expression and reversed fibrosis in patient skin explants[46]. Conservation across the four organs raises the possibility of broader relevance, but the producing cell, local activation mechanism and contribution relative to other TGF-β isoforms remain unresolved. Because systemic TGF-β interference can affect immune regulation and normal repair[1], isoform- and cell-resolved testing will be essential.

*ACKR1* points to the vascular-immune component of fibrosis. ACKR1-positive endothelial cells expand in human cirrhotic septa and enhance leukocyte transmigration[19]. In systemic sclerosis, ACKR1-high endothelium was linked to chemokine-producing stromal and immune populations, and experimental inhibition reduced leukocyte infiltration and fibrosis in bleomycin-treated mice[47]. The same role must be localised in heart, kidney and lung, and any blockade tested against the requirement for vascular homeostasis.

*AEBP1* and *FKBP10* act during matrix production. Aebp1 deficiency reduced myofibroblast accumulation and collagen deposition after bleomycin injury, while its collagen-associated product ACLP promoted fibroblast spreading, collagen contraction and myofibroblast differentiation[48]. Recombinant ACLP also increased smooth-muscle actin and collagen through TGF-β receptor-dependent and independent mechanisms[49]. FKBP10, an endoplasmic-reticulum collagen chaperone, was increased in idiopathic pulmonary fibrosis fibroblasts; knockdown reduced collagen I, collagen V, α-SMA and collagen secretion[50]. Their action before collagen enters mature scar makes perturbation studies particularly informative.

*GPR176* represents a distinct candidate class. This orphan G-protein-coupled receptor was increased across fibrotic mouse organs and concentrated in lung fibroblasts[51]. Knockdown in rat renal fibroblasts reduced TGF-β1-induced SMAD2 phosphorylation, α-SMA, fibronectin and collagen expression[51]. Evidence remains early because selective pharmacology and *in vivo* target blockade have not yet established a therapeutic effect.

*FBLN2* links the conserved signature to extracellular regulation of TGF-β. Loss of Fbln2 prevented angiotensin II-induced myocardial fibrosis despite comparable hypertension and hypertrophy, while recombinant fibulin-2 restored TGF-β signalling in deficient cardiac fibroblasts[52]. Whether this mechanism extends beyond the heart remains unresolved.

Most intervention evidence has been generated one target at a time, although fibrosis is maintained by interacting fibroblast, immune, endothelial and parenchymal programmes. Deleting TGF-β receptor 2 from matrix-producing renal interstitial cells reduced collagen I without reducing overall tissue fibrosis[53]. Dual MEK and PI3K inhibition reduced established TGF-β-driven pulmonary fibrosis more than either inhibitor alone[54], and concurrent LIGHT and TL1A blockade was more effective than either monotherapy after bleomycin injury[55].

These findings support testing combinations that cover complementary disease processes. Such combinations should span inflammatory recruitment, fibroblast activation and matrix maturation while preserving organ-specific repair and safety.

Direction is also critical. *EMILIN1* restrains extracellular maturation of pro-TGF-β, and its loss increased TGF-β signalling and caused late aortic-valve fibrosis[56, 57]. EFEMP2 supports elastic-fibre organisation and collagen assembly; loss reduced lysyl-oxidase activity and disrupted collagen cross-linking and fibril architecture[58]. Increased expression may therefore mark a compensatory repair response rather than a harmful dependency, so inhibition or activation should not be inferred from direction alone.

*OLFML2B*, *KDELR3*, *B3GNT9* and *SPECC1* have received little direct investigation in fibrosis. Their recurrence across all eight comparisons warrants cellular localisation and functional studies, but bulk data cannot distinguish causal regulation from expansion of a stromal population. They extend the candidate space beyond well-established matrix proteins.

The provisional biomarker also panel raises a related distinction. A recurrent tissue signature becomes a useful blood assay only if its products are released or shed, stable, measurable and sufficiently specific to separate fibrosis from inflammation, malignancy and normal turnover. Circulating MMP7 in idiopathic pulmonary fibrosis and PRO-C3 in liver disease demonstrate clinical potential, and a prespecified MMP7 prognostic threshold was reproduced when sampling procedures were comparable[59–61]. The ten-gene panel therefore requires assay development and external evaluation before any claim of clinical validity.

These findings must also be interpreted against heterogeneity in cohort composition, disease stage, sex balance, sequencing protocols and tissue handling. A condition-only DESeq2 design could not incorporate covariates consistently, and GSE213001 contains several anatomical regions from some donors. Human diseases and mouse injuries also differ in history and duration, while one effect-size threshold may favour larger or better-powered cohorts. Donor-aware reanalysis, cellular localisation, perturbation and independent cohorts are therefore required for confirmation.

In conclusion, fibrosis is therefore neither a uniform pan-organ programme nor a collection of wholly distinct tissue responses. A compact matrix-centred core sits within broader organ- and species-specific change. This distinction identifies candidates with potential relevance across organ-specific fibrotic diseases while preserving the need for tissue-specific validation. The next step is to determine which shared signals are causal and can be perturbed without disrupting essential repair or organ function.

## Data and code availability

All transcriptomic inputs were obtained from the public GEO, SRA and BioProject accessions listed in Table 1. ScarX V3.0 and its bundled demonstration datasets are available through the existing Hugging Face repository at https://huggingface.co/spaces/SamueleDiCarmine/scarx.

## Author contributions

**Conceptualization:** Samuele Di Carmine.

**Project administration:** Katie J. Mylonas.

**Resources:** Katie J. Mylonas.

**Visualization:** Samuele Di Carmine, Katie J. Mylonas.

**Writing – original draft:** Samuele Di Carmine.

**Writing – review & editing:** Richard Cunningham, Katie Mylonas.

