## Supplementary Table S1 for "A conserved matrix-centred core links fibrosis across human organs and mouse models"

**Supplementary Table S1. Twenty-five pan-organ gene-direction signatures conserved across human and mouse.**

| **Rank** | **Gene symbol** | **Cross-species definition** |
| --- | --- | --- |
| 1 | AEBP1 | Same gene and direction significant in all four human and all four mouse organs |
| 2 | B3GNT9 | Same gene and direction significant in all four human and all four mouse organs |
| 3 | CACNB3 | Same gene and direction significant in all four human and all four mouse organs |
| 4 | CDH11 | Same gene and direction significant in all four human and all four mouse organs |
| 5 | COL1A1 | Same gene and direction significant in all four human and all four mouse organs |
| 6 | COL1A2 | Same gene and direction significant in all four human and all four mouse organs |
| 7 | COL3A1 | Same gene and direction significant in all four human and all four mouse organs |
| 8 | COL5A2 | Same gene and direction significant in all four human and all four mouse organs |
| 9 | COL6A2 | Same gene and direction significant in all four human and all four mouse organs |
| 10 | DBN1 | Same gene and direction significant in all four human and all four mouse organs |
| 11 | EFEMP2 | Same gene and direction significant in all four human and all four mouse organs |
| 12 | EMILIN1 | Same gene and direction significant in all four human and all four mouse organs |
| 13 | FBLN2 | Same gene and direction significant in all four human and all four mouse organs |
| 14 | FKBP10 | Same gene and direction significant in all four human and all four mouse organs |
| 15 | FKBP4 | Same gene and direction significant in all four human and all four mouse organs |
| 16 | GPR176 | Same gene and direction significant in all four human and all four mouse organs |
| 17 | KDELR3 | Same gene and direction significant in all four human and all four mouse organs |
| 18 | LOXL1 | Same gene and direction significant in all four human and all four mouse organs |
| 19 | LOXL2 | Same gene and direction significant in all four human and all four mouse organs |
| 20 | OLFML2B | Same gene and direction significant in all four human and all four mouse organs |
| 21 | PMEPA1 | Same gene and direction significant in all four human and all four mouse organs |
| 22 | RAB7B | Same gene and direction significant in all four human and all four mouse organs |
| 23 | SPECC1 | Same gene and direction significant in all four human and all four mouse organs |
| 24 | TGFB2 | Same gene and direction significant in all four human and all four mouse organs |
| 25 | VIM | Same gene and direction significant in all four human and all four mouse organs |
