## Supplementary Table S2 for "A conserved matrix-centred core links fibrosis across human organs and mouse models"

**Supplementary Table S2. Local tractability display classes among the top 500 HCM-ranked candidates.**

| **Local tractability display class** | **Top-500 count** | **Percentage** |
| --- | --- | --- |
| Cell surface | 153 | 30.6% |
| Cell surface and kinase | 8 | 1.6% |
| Kinase | 1 | 0.2% |
| Secreted | 133 | 26.6% |
| Secreted and cell surface | 64 | 12.8% |
| Transcription factor | 17 | 3.4% |
| No local tractability class | 124 | 24.8% |
