## Supplementary Figure S1 for "A conserved matrix-centred core links fibrosis across human organs and mouse models"

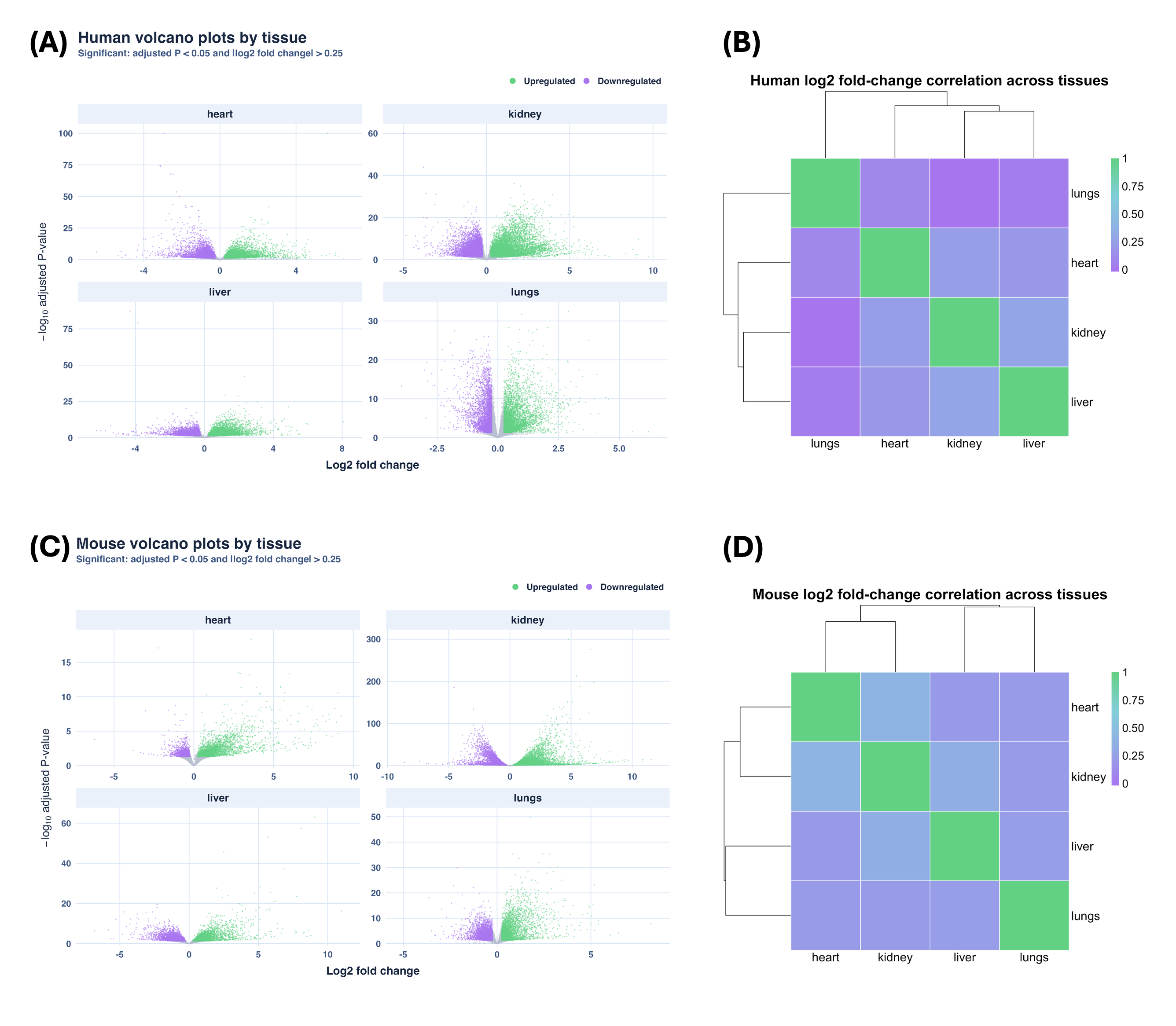

Supplementary Figure S1. Organ-level differential-expression distributions and within-species effect correlations. Volcano panels display apeglm-shrunken log2 fold change against -log10 adjusted P for human (A) and mouse (C) organ comparisons; thresholds were adjusted P < 0.05 and absolute shrunken log2 fold change > 0.20. Heatmaps show pairwise correlations among complete organ-level log2 fold-change profiles in human (B) and mouse (D).
