## Supplementary Figure S2 for "A conserved matrix-centred core links fibrosis across human organs and mouse models"

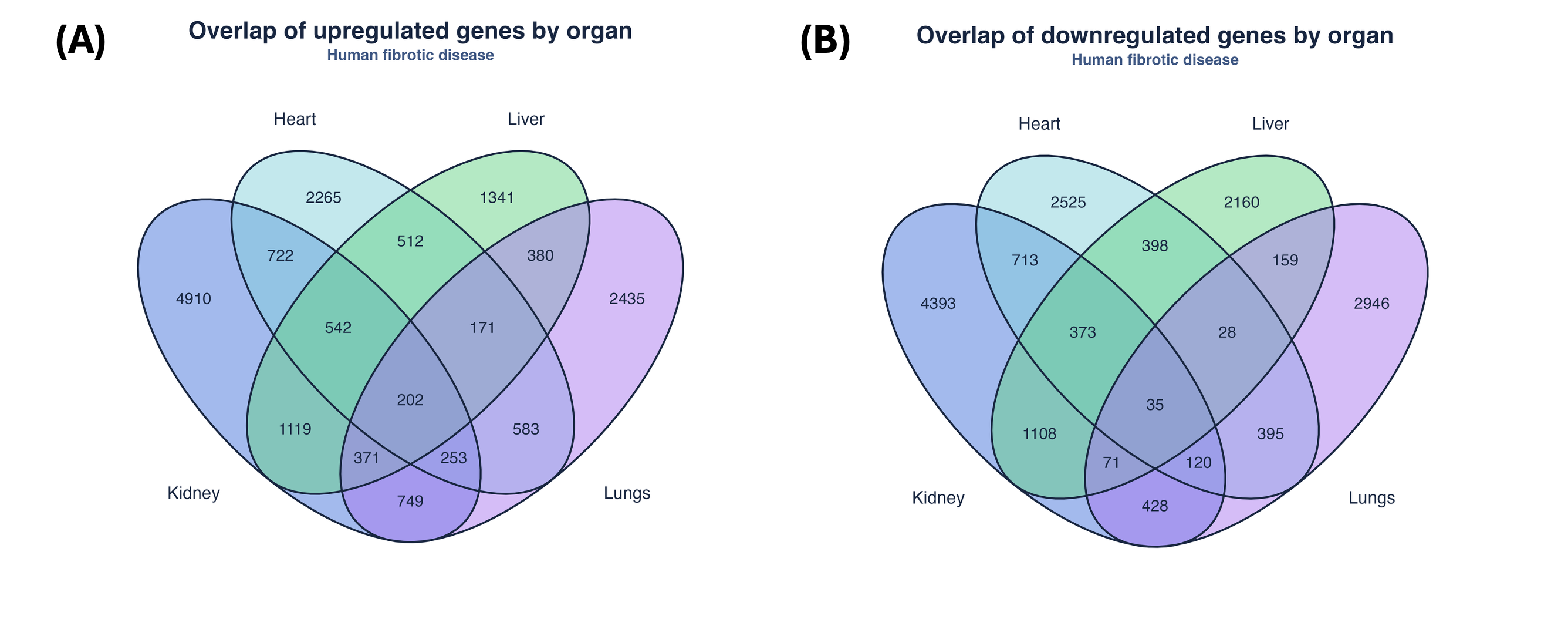

Supplementary Figure S2. Human cross-organ set overlap. Venn diagrams show the intersections of significantly increased (A) and decreased (B) genes among heart, kidney, liver and lung. Significance required adjusted P < 0.05 and absolute apeglm-shrunken log2 fold change > 0.20.
