## Supplementary Figure S3 for "A conserved matrix-centred core links fibrosis across human organs and mouse models"

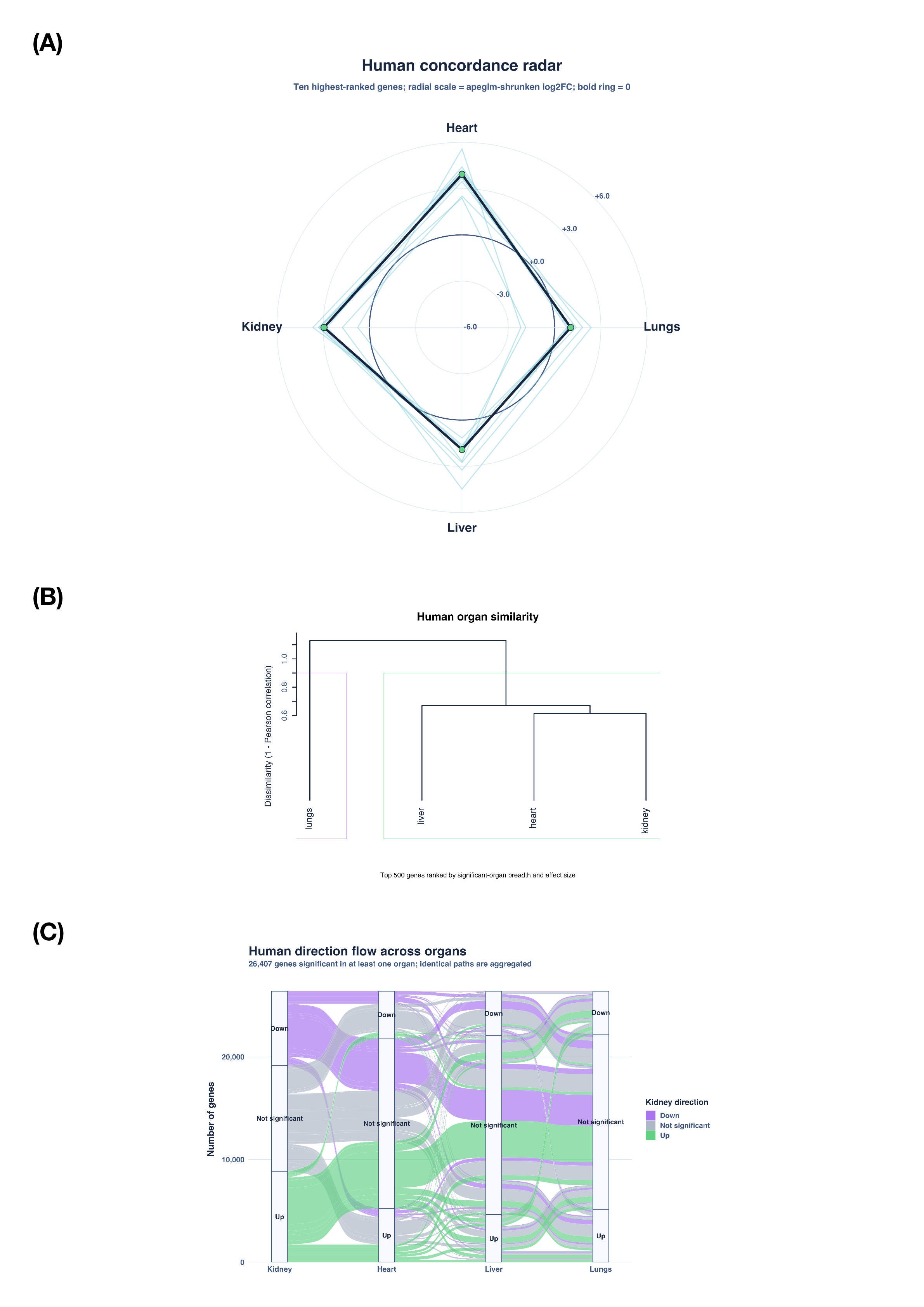
 Supplementary Figure S3. Exploratory structure of the human cross-organ output. (A) Signed apeglm-shrunken log2 fold-change profiles for the ten highest-ranked human genes. Genes were pre-ranked by the number of organs meeting the significance criteria, mean absolute effect across significant organs and maximum absolute effect; the 500 highest-ranked genes were then ordered by mean absolute effect across all four organs. Pale lines show individual genes and the bold line shows the organ-wise median. The signed display scale spans -6 to +6 apeglm-shrunken log2 fold change; no displayed value was clipped. (B) Hierarchical clustering of organs based on log2 fold-change correlation. (C) River plot showing transitions among up, down and non-significant classes. Significance required adjusted P < 0.05 and absolute shrunken log2 fold change > 0.20.
