## Supplementary Figure S5 for "A conserved matrix-centred core links fibrosis across human organs and mouse models"

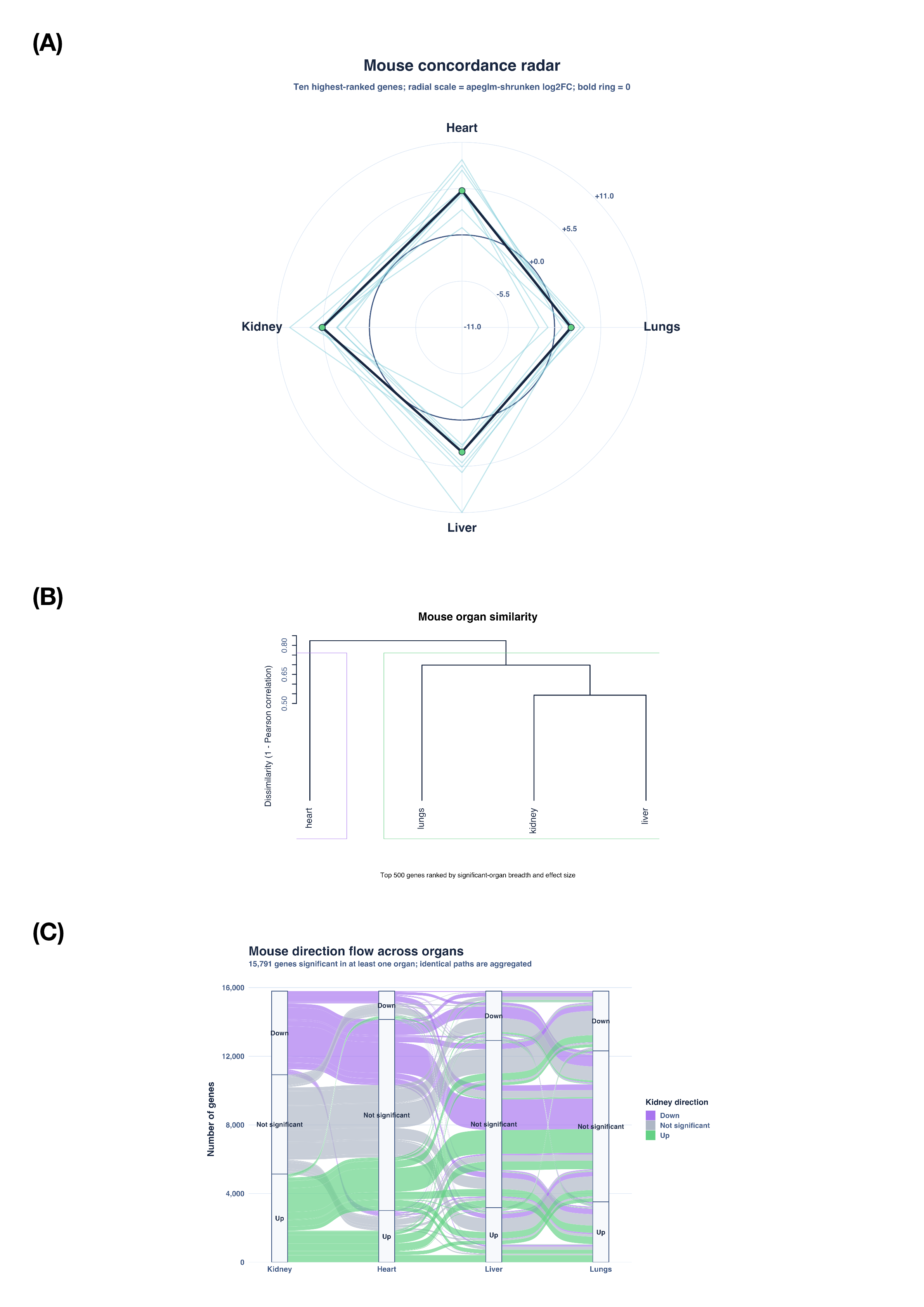
 Supplementary Figure S5. Exploratory structure of the mouse cross-organ output. (A) Signed apeglm-shrunken log2 fold-change profiles for the ten highest-ranked mouse genes, selected using the same criteria as in Supplementary Figure S3A. Pale lines show individual genes and the bold line shows the organ-wise median. The signed display scale spans -11 to +11 apeglm-shrunken log2 fold change; no displayed value was clipped. (B) Hierarchical clustering of organs based on log2 fold-change correlation. (C) River plot showing transitions among up, down and non-significant classes Significance required adjusted P < 0.05 and absolute shrunken log2 fold change > 0.20.
