## Supplementary Figure S6 for "A conserved matrix-centred core links fibrosis across human organs and mouse models"

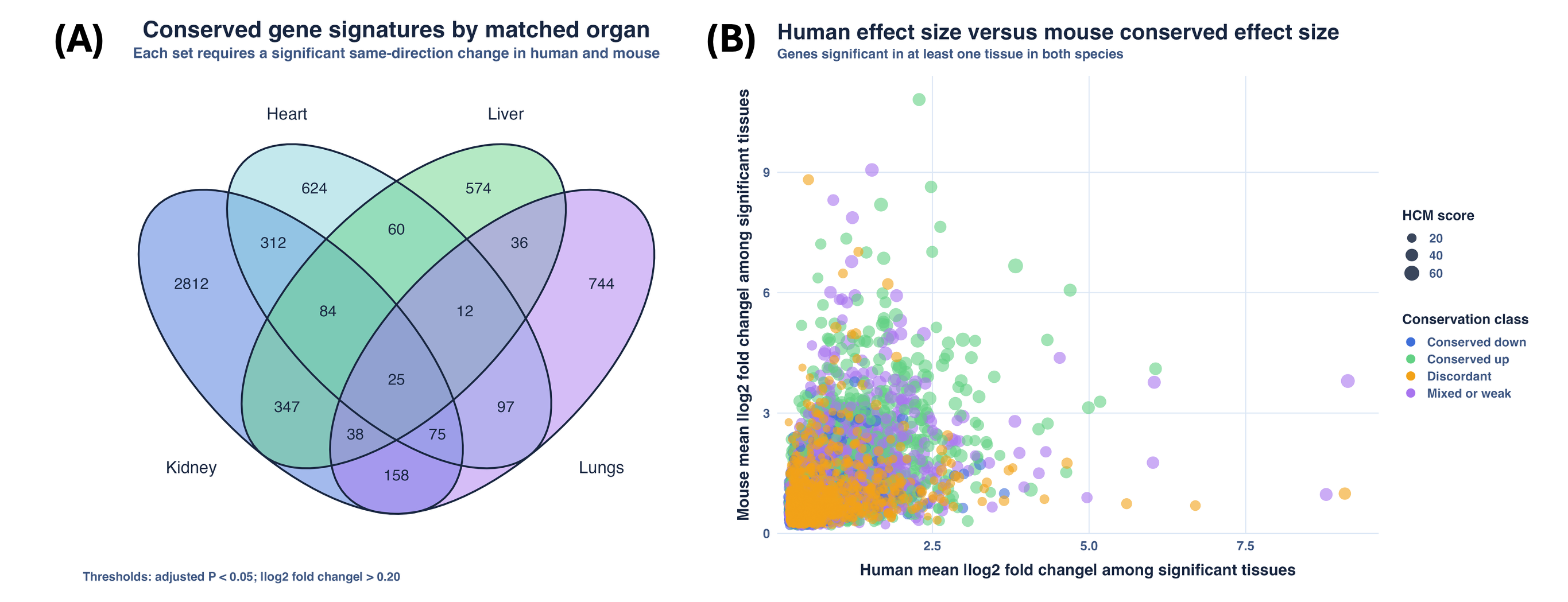

Supplementary Figure S6. Organ breadth and cross-species effect distribution. (A) Four-set Venn diagram of genes significant in the same direction in human and mouse within each matched organ. (B) Human mean absolute apeglm-shrunken log2 fold change plotted against mouse mean absolute apeglm-shrunken log2 fold change for matched genes; point colour shows conservation class and point size shows HCM score.
